# Imaging the structural connectome with hybrid diffusion MRI-microscopy tractography

**DOI:** 10.1101/2024.01.08.574641

**Authors:** Silei Zhu, Istvan N. Huszar, Michiel Cottaar, Greg Daubney, Nicole Eichert, Taylor Hanayik, Alexandre A. Khrapitchev, Rogier B. Mars, Jeroen Mollink, Jerome Sallet, Connor Scott, Adele Smart, Saad Jbabdi, Karla L. Miller, Amy F.D. Howard

## Abstract

Neuroanatomical tract tracing methods are fundamental in providing “gold standard” estimates of brain connectivity. However, tracer methods cannot be performed in humans and even in animals, we can only study projections from typically one or two injection sites per animal sacrificed. Orientation-sensitive microscopy techniques such as PLI provide an alternative where they can visualise detailed fibre orientations at the micron-scale across the whole brain. However, these methods are often most informative on orientations within the 2D imaging plane, with less reliable or missing through-plane information, restricting 3D tract reconstruction. Conversely, dMRI can estimate fibre orientations in 3D but at low resolution, which leads to many false positive and negative estimates of fibre trajectories.

To facilitate reconstruction of the microscopy-informed connectome, we develop a data-fusion method that complements 2D microscopy with through-plane information from diffusion MRI to construct 3D hybrid orientations that are both maximally informed by the high-resolution microscopy, have whole-brain coverage and can be input into existing tractography pipelines. Diffusion MRI can be readily acquired prior to microscopy meaning the same method is translatable across species, including in humans. Here we apply our method to an existing open-access macaque dataset and demonstrate (1) whole-brain microscopy-informed tractography (2) the advantages of hybrid tractography in two known tractography challenges, the gyral bias and bottleneck problem (3) how hybrid tractography appears to outperform diffusion-only tractography when compared to tracer data and (4) the generalisability of our hybrid method to different microscopy contrasts, facilitating wider translation.

## 2. Introduction

Mapping brain structure through multiple modalities can improve our understanding of brain function, development, aging and diseases^1–4^. Much effort has gone into creating comprehensive maps of brain connectivity in different species using various methods from MRI to microscopy^5–7^. Different modalities can provide complementary information about the underlying tissue microstructure by probing different length scales and providing sensitivity to different tissue features. Notably, when multiple modalities are combined in the same tissue, this can offer new opportunities for multi-scale neuroscience.

Chemical tracers have been an important tool for mapping the pathways of neural connections in animal brains over many years^8–11^. The tracer facilitates direct visualisation of axonal projection from an injection site, providing a highly specific and sensitive estimate of inter-regional brain connectivity^11,12^. However, tracers are limited to a single or a few injections per animal, where achieving whole-brain coverage requires sacrificing many animals, and combining information across subjects ignores important between-subject variability^8,11,12^. Further, tracers are not available in humans.

Alternative microscopy techniques serve as crucial reference measurements for characterising fibre architecture at high spatial resolution^13^. For microscopy to be useful in tract reconstruction, the microscopy must inform on the fibre orientations in 3D. Though multiple microscopy methods can provide 3D information (e.g. 3D polarised light imaging [3D-PLI]^14–16^, μ-CT^17–19^, SAXS^20^ or serial EM^21^), these typically require more sophisticated hardware and long acquisition times which can limit widespread application. Consequently, microscopy orientations are often acquired in 3D from only small tissue samples or in 2D from thin sections of brain tissue, thus precluding 3D tract reconstruction or whole-brain connectivity estimates in larger (e.g. primate) brains.

In comparison, diffusion MRI (dMRI) enables non-invasive mapping of structural brain connectivity^22^. By measuring the diffusive motion of water molecules through the tissue, we can infer underlying fibre orientations and reconstruct white matter (WM) fibre bundles or tracts via tractography methods^23^. This facilitates the whole-brain estimation of white matter organisation but relies on fibre orientations estimated via computational models from millimetre-scale MR signals. Low resolution and inaccuracies in the models can introduce bias or noise in the inferred fibre orientations that contribute to false positives and negatives in downstream tractography outputs^24–26^.

Here, we propose a data-fusion approach to jointly analyse data from dMRI and microscopy to perform whole-brain microscopy-informed tractography. Our framework leverages the complementary information these modalities provide to create hybrid dMRI-microscopy fibre orientations that are both 3D and at high resolution (Figure 1), preserving the unique benefits of the microscopy whilst facilitating 3D tractography. We use 2D microscopy to provide detailed estimates of fibre orientation within the microscopy plane and dMRI to provide the through-plane orientation. For the latter, we use models that estimate distributions of fibre orientations within dMRI voxels^27^, rather than a single orientation per voxel (e.g. from DTI)^28^. By extracting the through-plane angle for the fibre within this distribution that best matches the in-plane orientation derived from microscopy, we can then estimate 3D fibre orientations at spatial resolutions that exceed the dMRI data. Our hybrid dMRI-microscopy method provides: 1) 3D fibre orientations; 2) whole-brain coverage; 3) high resolution information and estimation of complex fibre architecture as described by the microscopy. The hybrid orientations are then combined into fibre orientation distributions (FODs) at arbitrary resolutions that are input into existing tractography pipelines for tract reconstruction. When based on myelin-sensitive microscopy, the hybrid orientations likely provide a more “myelin-specific” fibre orientation distribution than dMRI, where the dMRI FOD can represent a range of microstructural features including axons, dendrites and glia processes. These myelin-specific FODs may be advantageous for certain applications such as defining the “myelin connectome” or tracking myelinated fibres into the cortex.

**Figure 1:**
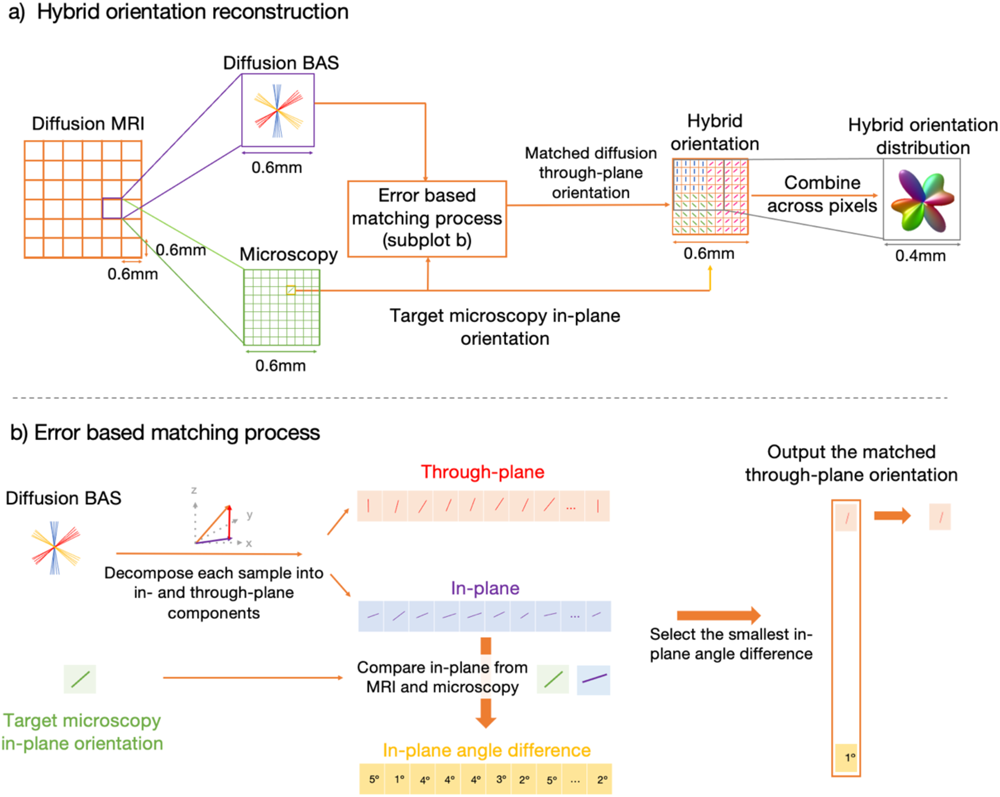
Overview of the hybrid orientation approach. The hybrid orientation method combines microscopy and dMRI for high-resolution 3D fibre orientation. a) The dMRI data was analysed with the ball and stick model (BAS) at 0.6mm. Each microscopy pixel and the diffusion BAS within the same voxel were used in an error-based matching process. This process calculated the matched diffusion through-plane orientation, which was combined with the 2D microscopy orientation to reconstruct the hybrid orientation. This approach provides microscopic resolution and allows combining orientations into a FOD represented by spherical harmonics (Supplementary Figure 1, Supplementary Figure 2). These FODs can then be input into existing tractography algorithms for investigations of whole-brain structural connectivity (Supplementary Figure 3). b) Error-based matching process. The diffusion BAS and microscopy in-plane orientation were used as inputs. The 3D BAS orientation was decomposed into the through-plane angle and in-plane angle by projecting onto the microscopy plane. This process was repeated for multiple fibre populations (≤ 3) with 50 orientation samples per population of the BAS. The in-plane angle was compared to the target microscopy in-plane orientation by quantifying the angle difference. Supplementary Figure 4 shows the in-plane, through-plane angle and angle difference maps. We selected the BAS sample with the smallest in-plane angle difference and determined the matched through-plane angle. Thus, the BAS orientation samples reflected the distribution of fibre orientations throughout the dMRI voxel. However, the precise sub-voxel localisation of these fibre orientations was unknown. The microscopy essentially assigned the through-plane orientation of the 3D samples to their putative location within the voxel.

We demonstrate our approach using the BigMac dataset^29^, an open-access multi-modal resource that includes postmortem dMRI and microscopy data from a single macaque brain with whole-brain coverage. The microscopy includes polarised light imaging (PLI)^25–27^, myelin^33^ and Nissl-stained^34,35^ histology. Precise co-registration between microscopy and MRI has been conducted, which is essential for meaningful data fusion at the voxel-wise level^36^. Applying our hybrid method to the dMRI and PLI data, we first demonstrate how to perform hybrid tractography at different resolutions and reconstruct 42 microscopy-informed tracts spanning the whole macaque brain. With confidence in our method, we then demonstrate how hybrid tractography can be beneficial in neuroanatomical and methodological investigations. Specifically, we utilise our hybrid outputs to investigate two known challenges in dMRI-based tractography: the gyral bias and bottleneck problem (Figure 2), both of which are primarily driven by the limited spatial resolution of dMRI data. The gyral bias describes an established preference for dMRI fibre trajectories (streamlines) to terminate at gyral crowns rather than accurately capturing sharp turns into gyral walls^24,25^. In bottleneck regions, multiple fibre bundles merge causing streamlines to become indistinguishably mixed at the spatial resolution of dMRI^26,37^. Without additional anatomical knowledge, it is difficult to retain information about the origin and termination of individual tracts passing through a bottleneck. By comparing our hybrid outputs to dMRI results in the same brain, we investigate how these challenges are related to the resolution and contrast-generating mechanism of dMRI. We then compare the hybrid tractography outputs to tracer data obtained from other animals^11,12^ to demonstrate that our hybrid method provides higher specificity in fibre tracking than diffusion only tractography. Finally, we perform hybrid tractography using three different microscopy contrasts (PLI, myelin-and Nissl-stained histology) to demonstrate its generalizability. Overall, our method retains the benefits of microscopy without relying on invasive tracers, meaning we can estimate dense, microscopy-informed structural connectivity from a single brain, using a method that is translatable across species, including humans.

**Figure 2:**
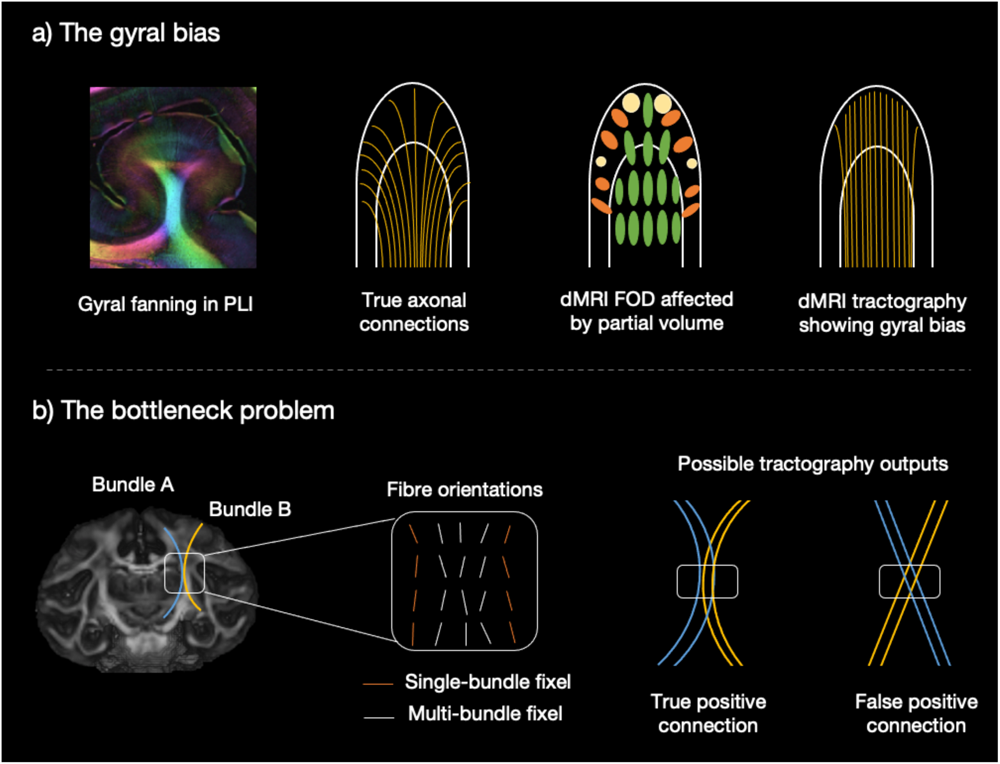
The tractography challenges. a) The ground truth gyral fanning is demonstrated using PLI and a schematic representation of fibre connections. However, in dMRI, the accuracy of FODs is affected by the partial volume effect. The estimated diffusion tractography trajectories (streamlines) predominantly terminate at the gyral crowns but fail to turn into the gyral wall, resulting in the gyral bias problem. b) A bottleneck example in the internal capsule is shown (Adapted from Schilling et al., 2022). Two fibre bundles, originating and terminating at different locations converge with similar orientations within the bottleneck region (truth axonal connection). Fixels are classified based on the number of fibre bundles passing through each fixel: multi-bundle fixels (white) have multiple associated fibre bundles and single-bundle fixels have a single associated fibre bundle (orange). The bottleneck region, identified as the multi-bundle fixel where streamlines become indistinguishably mixed, generates invalid streamlines (false positive connection). As a result, a single FOD pattern can produce two probabilistic tractography outputs.

## 3. Results

### 3.1 Generating hybrid dMRI-PLI fibre orientations

Hybrid dMRI-microscopy orientations were estimated for each microscopy pixel, where microscopy provided the in-plane orientation and dMRI approximated the through-plane orientation. The through-plane orientation was determined with an error-based matching process described in Figure 1. The dMRI data was first analysed using the Ball and Stick (BAS) model^27^ to estimate up to 3 fibre populations per voxel, with 50 orientation estimates or ‘samples’ per population. These sample orientations capture the uncertainty in the estimation of the 3 fibre populations, which in part reflects the spatial distribution of the fibres across the voxel. The 2D microscopy was then used to spatially assign the through-plane orientation of these fibres to the high-resolution sub-voxel grid of the microscopy. Each 2D microscopy orientation was warped to dMRI space and compared to the BAS samples in the same voxel, where the BAS samples were first projected onto the microscopy plane to facilitate fair comparison with the 2D microscopy. The microscopy through-plane information was then approximated using that from the most similar BAS sample. This approach yielded 3D fibre orientations at the resolution of the microscopy. An additional benefit is that by utilising a dMRI fibre reconstruction method that produces multiple fibre estimates per voxel, we have the opportunity to “super-resolve” 3D fibre orientations at a resolution higher than the native MRI. We use the term dMRI-microscopy to refer to the general method, as multiple microscopy contrasts can be used to create hybrid orientations. dMRI-PLI is used to denote the hybrid orientations reconstructed using PLI which estimates the in-plane fibre orientation based on tissue birefringence^16,30–32^.

The hybrid orientations can then be combined over a local neighbourhood to create fibre orientation distributions (FODs, here described using spherical harmonics) that are directly comparable to FODs output from dMRI, and can be input into existing tractography algorithms for tract reconstruction^38^. One important feature of our hybrid method is that the hybrid FODs can be constructed at different resolutions from the same underlying data by changing the size of the combined neighbourhood region (Supplementary Figure 1). This allows us to investigate how spatial resolution affects tractography without confounding factors such as signal-to-noise ratio (SNR) changes in MRI acquired at different resolutions. Supplementary Figure 2 demonstrates how using 0.6mm resolution postmortem dMRI and PLI microscopy at 4 μm/pixel we created hybrid FODs at multiple resolutions: 1mm isotropic, 0.6mm isotropic, 0.4mm isotropic, and 0.1x0.4x0.1mm (In the latter, the through-plane resolution (0.4mm) was determined by the distance between consecutive PLI slices (0.35mm).

The hybrid FODs at 0.6mm isotropic were compared to the orientations obtained from the dMRI data using both ball and stick model (BAS) and constrained spherical deconvolution (CSD) approaches (Figure 3). Notably, the CSD output also uses spherical harmonics making the FODs easy to compare. For the hybrid FODs, the microscopy mostly informed the orientations within the coronal plane, whilst the dMRI provided through-plane information in the sagittal and transverse views. We observe how the hybrid dMRI-PLI FODs agree with the fibre orientations from BAS and CSD in all three views and faithfully depict U-shaped fibres connecting cortical areas between adjacent gyri. Together, this gives confidence in the fidelity of the hybrid outputs.

**Figure 3:**
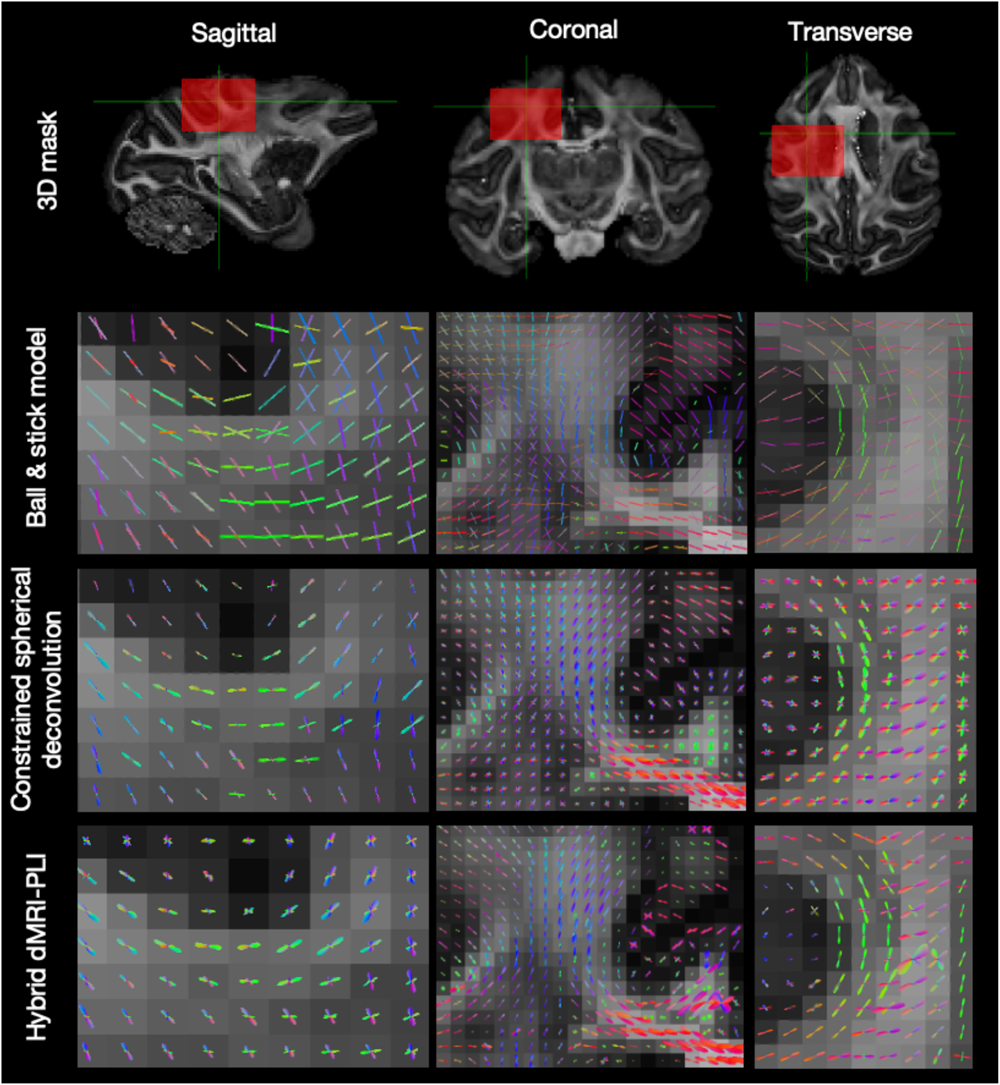
Comparison between dMRI FOD and hybrid dMRI-PLI FOD. The FOD generated from ball and stick model (Top), constrained spherical deconvolution (Middle) and hybrid dMRI-PLI (Bottom). The orientations in sagittal, coronal and transverse views are presented for a region near to the grey matter.

### 3.2 Whole-brain hybrid dMRI-PLI tractography

Using fibre orientation estimates from the hybrid dMRI-PLI, fibre bundles were reconstructed with a standard tractography pipeline (Supplementary Figure 3D). Figure 4 displays the reconstruction of three example tracts at the hybrid spatial resolutions of 0.4mm, 0.6mm, and 1mm isotropic, alongside tractography using conventional dMRI-only analysis (constrained spherical deconvolution) at 0.6mm. We specifically chose tracts that represent two extremes with respect to how the hybrid orientations relate to the underlying data: (i) the cortico-spinal tract (CST) predominantly runs in parallel to the microscopy plane such that microscopy is most informative; (ii) the optic radiation extends through the microscopy plane for which dMRI is most informative. Hybrid and dMRI tractography produce similar reconstructions of both tracts. Both tracts are characterized by intricate anatomical features: the CST tracks through multiple crossing fibre regions, with branches extending to both superior and lateral regions of the motor cortex; the optic radiation contains a backward curving structure known as Meyer’s Loop in the anterior portion^39^. The hybrid method successfully reconstructs both complex structures. The uncinate fibre tract represents a case where the hybrid tractography does not perform so well. Most of the uncinate fibre can be robustly reconstructed with a clearly observed U shape structure extending from the anterior temporal lobe to the orbitofrontal cortex. However, the hybrid method fails to capture the most anterior part of the uncinate tracking through the orbitofrontal cortex, which is well reconstructed using dMRI-tractography. The limited anterior extent may be caused by registration inaccuracies in the most anterior part of the brain due to the limited amount of brain tissue and less distinct anatomical features to drive the registration.

**Figure 4:**
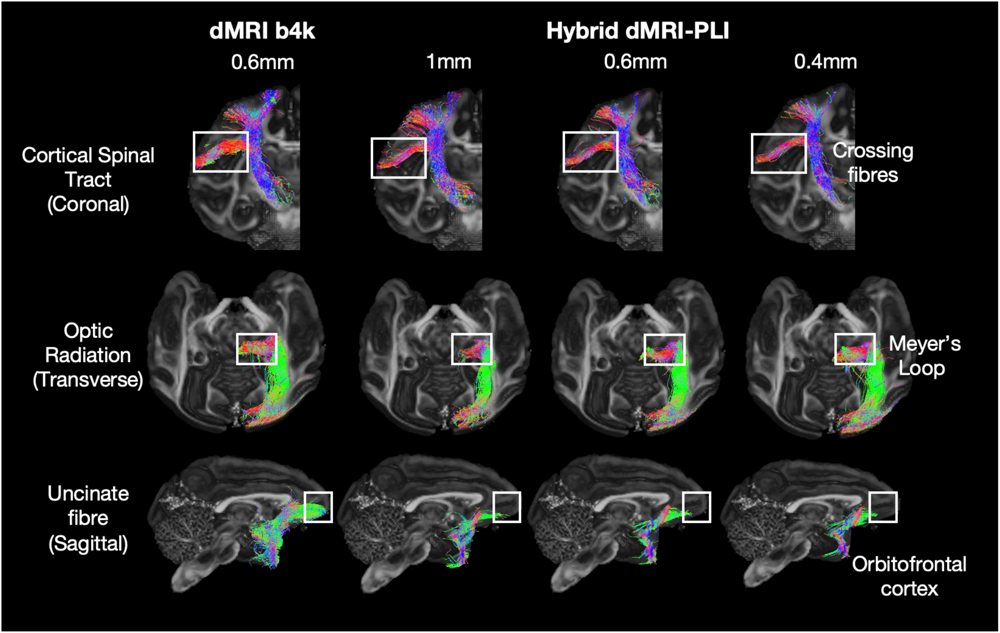
Example tracts generated using the hybrid tractography. The hybrid method can successfully reconstruct numerous tracts throughout the brain including the CST, an example tract within the coronal plane where microscopy is the most informative (Top), the optic radiation, a tract extending primarily along the anterior-posterior axis where dMRI provides most information (Middle), and the uncinate fibre (Bottom), a tract connecting the anterior temporal lobe and the orbitofrontal cortex.

Having established that our method can provide reliable tract reconstructions, we then proceeded to reconstruct fibre bundles spanning the whole brain (Supplementary Figure 6). The XTRACT toolbox was employed to define seed and target regions of interest (ROIs) for a total of 42 tracts, comprising association, commissural, and projection fibres^40^.

The BigMac macaque brain was found to have a cerebral bleed in the right hemisphere, as observed in the MRI (loss of signal) and microscopy (missing tissue). This bleeding site primarily affected the grey matter but had some overlap with tracts including SLF2. Consequently, we observed asymmetry in the reconstruction of SLF2 between the left and right hemispheres (Supplementary Figure 7), where streamlines failed to track through the bleeding site on the right. Despite this limitation, other tracts were successfully reconstructed using the hybrid orientations, yielding anatomically expected structures (Figure 5). The results further demonstrate that our method is applicable to a wide range of major white matter tracts and can reconstruct tracts spanning the whole macaque brain.

**Figure 5:**
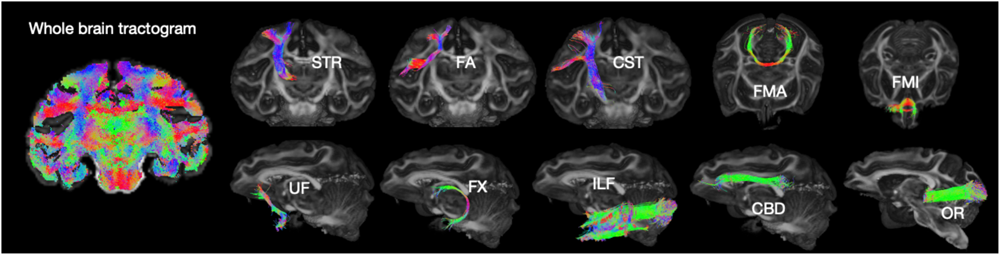
Hybrid dMRI-PLI tracts spanning the whole brain. Whole-brain tractogram is shown. Ten example tracts generated from the hybrid tractography at 0.6mm isotropic are illustrated including the superior thalamic radiation (STR), frontal aslant (FA), corticospinal tract (CST) in the coronal view, forceps major (FMA), forceps minor (FMI) in the axial view, uncinate fasciculus (UF), fornix (FX), inferior longitudinal fasciculus (ILF), cingulum subsection: dorsal (CBD), optic radiation (OR) in the sagittal view.

Although the hybrid method accurately captures the course of the uncinate fibre, it shows less anterior extent than dMRI-based reconstruction. The anatomical features of interest are labelled with white boxes.

### 3.3 Hybrid tractography eliminates the gyral bias at high resolution

At the spatial resolution available to dMRI scans, the signal in voxels near the cortex is dominated by fibres projecting towards the gyral crowns. This causes streamlines to run parallel to the gyral walls until they reach the crown, leading to biases in connectivity mapping. This so-called gyral bias is in part due to the spatial resolution of dMRI^24,41^. Our hybrid method provides microscopy-informed fibre orientations that can be reconstructed at multiple spatial resolutions down to the near-native microscopy resolution. We investigated whether this approach can reduce or eliminate the gyral bias problem. In Figure 6, the gyral bias is evident in the 1mm hybrid data, with fibres preferentially terminating at the gyral crown. However, with increased spatial resolution we observe an increasing number of streamlines extending towards the gyral walls, aligned with previous observations from microscopy^41^. The dMRI-only data shows similar trends of reduced gyral bias with increased spatial resolution, though with a less clean fanning pattern than that observed in the hybrid results. These results suggest that increasing the spatial resolution in both diffusion and hybrid tractography can overcome the gyral bias problem which is consistent with previous findings^24,41^.

**Figure 6:**
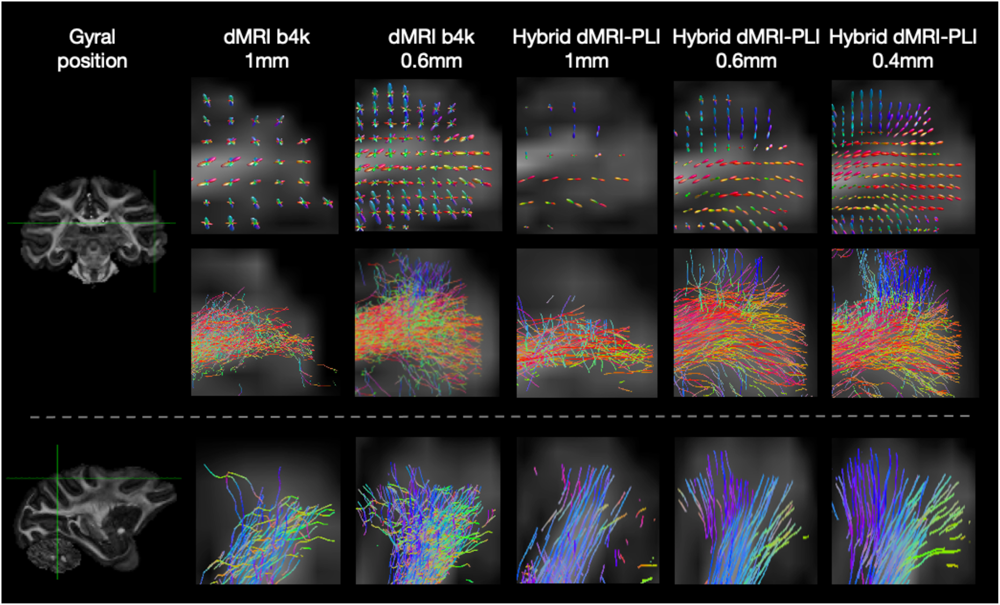
High resolution hybrid tractography eliminates the gyral bias. FODs (Top) and tractography streamlines of two gyri which primarily lie within (Middle) and through (Bottom) the microscopy plane. Outputs are shown for dMRI and hybrid dMRI-PLI reconstructed at different isotropic resolutions. The gyral bias problem is present in the 1mm, whilst the high spatial resolution of 0.4mm successfully delineates the expected fibre fanning.

### 3.4 Hybrid tractography preserve topography in bottleneck areas

Another limitation of dMRI tractography is the so-called bottleneck problem. When white matter pathways funnel through a single voxel, this can cause streamlines from different pathways to become intertwined and lose their topographic arrangement. This problem is exacerbated by the low resolution of dMRI relative to white matter axons. We further investigated the use of hybrid method in tackling bottleneck regions to ask whether spatial resolution and/or the microscopy-informed fibre orientations can provide superior tract separation than typical dMRI analysis. Our first bottleneck of interest was the internal capsule (IC). This was motivated by the known topographic arrangement of fibres within this tract, mirroring the topography of the motor cortex. The medial-to-lateral functional organisation of the motor cortex has a topographic organisation, which is maintained along the anterior-to-posterior axis of the IC, with fibres originating from medial “trunk (and legs)” regions located anteriorly within the IC compared to those from lateral “face” regions^42^. Streamlines were seeded from three ROIs defined on the precentral gyrus representing the trunk (and legs), arm and face regions with a medial-to-lateral distribution (Figure 7a). Figure 7b shows voxels in the IC colour-coded by the ROI with the highest streamline density. Figure 7c visualises the streamlines from each ROI projecting through the IC. Whilst dMRI mixes streamlines from distinct ROIs as they pass through the IC, the hybrid method consistently retains the strong topographic organisation of streamlines through the bottleneck region, where the order of trunk-arm-face regions along the anterior-posterior axis in the IC follows neuroanatomical expectations^42^.

**Figure 7:**
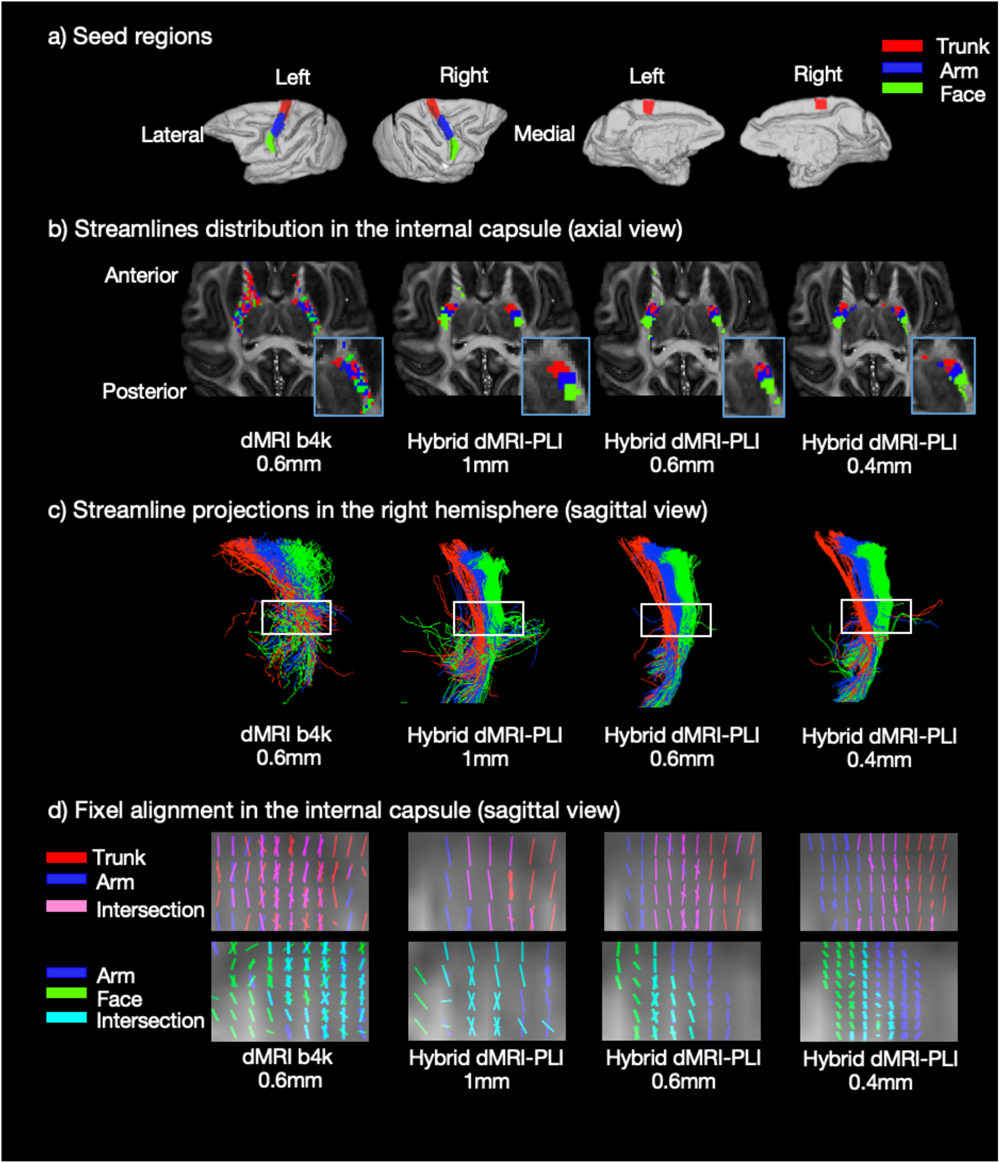
Hybrid tractography preserves topography in the internal capsule. a) ROIs relate to the functional representation of the trunk, arm and face regions shown for both medial and lateral parts of the two hemispheres. Tractography was seeded from the ROIs to reconstruct streamlines passing through the internal capsule (IC). (b) The density map and (c) streamline projections are shown. The blue box indicates a zoomed region in (b) and the white box in (c) is the bottleneck region of interest. The bottleneck problem is observed in the dMRI as streamlines from the ROIs are mixed. With the hybrid method, the streamlines from each ROI demonstrate a clear anterior-posterior distribution in the bottleneck region. Notably, the ROIs are separable at all hybrid resolutions. d) Fixel-based analysis was performed to generate a fixel density map for each ROI in the internal capsule. The red, blue and green colours represent fixels associated with the trunk, arm and face regions respectively. Pink shows fixels associated with both the trunk (red) and arm (blue) ROIs. Cyan shows the intersection of fixels from both the arm (blue) and face (green).

Fixel-based analysis was then used to visualise the fibre populations in each voxel associated with each tract^43^, quantifying tract orientation overlap within the bottleneck (Supplementary Figure 8). In each voxel, the FOD was divided into discrete fibre populations, or “fixels”. During tractography, each streamline passed through a voxel by following a specific fixel. Consequently, we can describe each streamline as a series of fixels projecting from our seed ROI. Combining across all streamlines in a given tract produced a fixel density map. This is analogous to a streamline density map, but now on a fixel rather than voxel basis. The density map was then converted into a tract-specific fixel mask using a 5% threshold.

Regions where a single fixel is transversed by multiple fibre bundles (referred to as a “multi-bundle fixel”) are defined as bottleneck regions. Conversely, fixel associated with a single bundle is termed “single-bundle fixel”^26^. Single-bundle fixels are colour coded using red, blue and green according to the ROI from which they are seeded (red=trunk, blue=arm, green=face). Multi-bundle fixels exhibit overlapping colours such as pink resulting from the overlap between the red (trunk) and blue (arm) fixels, or cyan resulting from the overlap between blue (arm) and green (face) fixels. In the dMRI analysis, we observe a large number of multi-bundle fixels (i.e. a large bottleneck region). This suggests that streamlines originating from different motor cortex ROIs tend to jump from one tract to the other. In the hybrid results, there is a clear separation of the ROIs on either side, with an intersecting bottleneck region in the middle where the two pathways run parallel to each other with shared orientations. This pattern is most evident in the 0.4mm, as with increased spatial resolution, the bottleneck region decreases in size and the separation becomes clearer.

Together, the results demonstrate how, irrespective of the spatial resolution, microscopy-informed tractography mitigates the bottleneck problem in the IC. Interestingly, this holds even for the 1 mm hybrid reconstruction, a coarser resolution than that of the dMRI data used in this comparison (0.6mm).

A second bottleneck region in the brainstem was studied using tractography seeded from ROIs in the primary motor and somatosensory cortex (Figure 8)^44^. As above, density maps and tractography results are presented for dMRI and hybrid orientation at different spatial resolutions. The bottleneck problem is observed in both the dMRI and the hybrid method at 1mm as streamlines from the two ROIs are mixed. In the hybrid high-resolution results (0.4mm and 0.6mm), the streamlines from each ROI demonstrate a separable distribution. Streamlines from the primary motor ROI are observed to be more anterior to those from the somatosensory ROI, mirroring the anterior-posterior topographic organisation of the cortex. As in the IC, the fixel-based analysis indicates how at higher spatial resolution there are fewer fixels associated with the bottleneck (pink), facilitating better tract separation.

**Figure 8:**
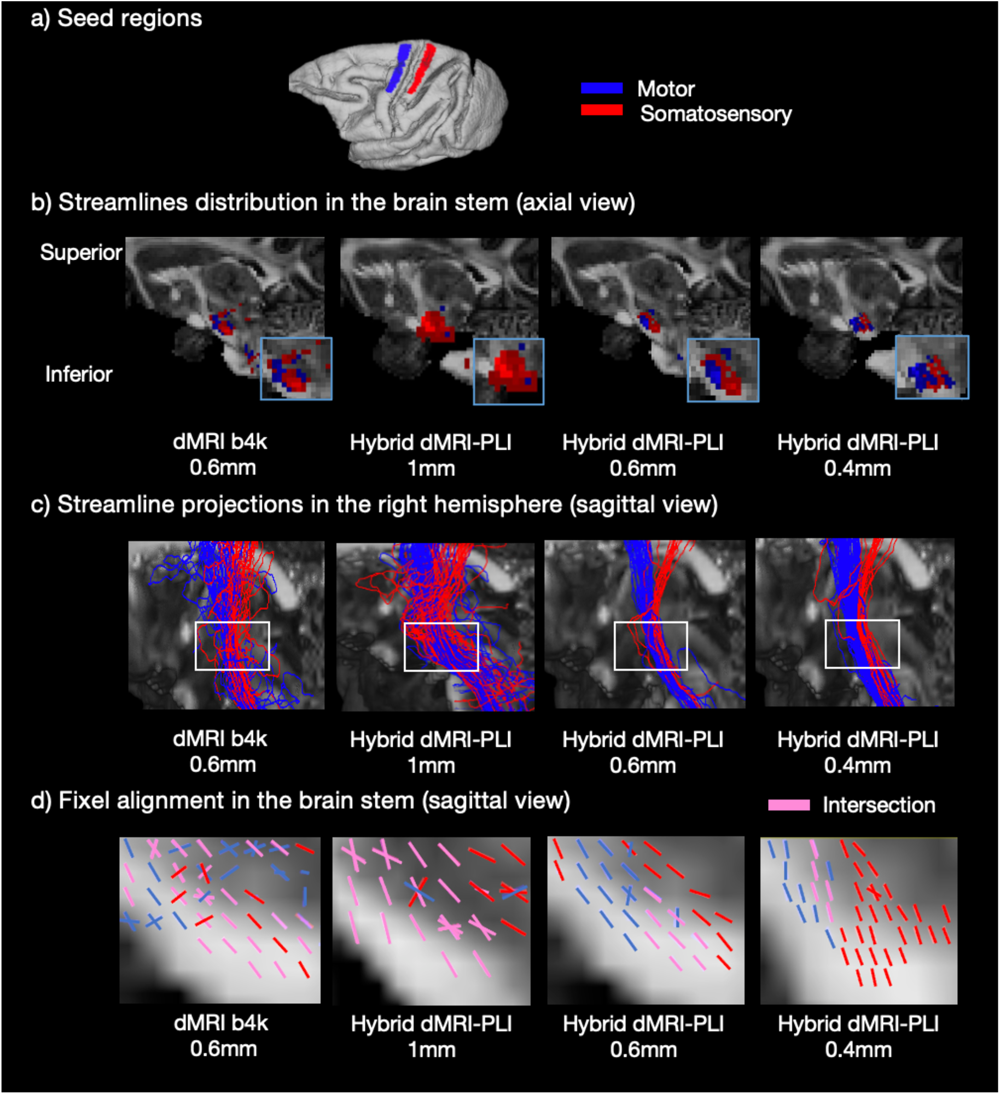
Hybrid tractography preserves topography in the brainstem. a) ROIs in the primary motor (blue) and somatosensory (red) cortex. Tractography was seeded from the ROIs to reconstruct streamlines passing through the brainstem. b) The density map in sagittal view and (c) the streamline projections are shown. The blue box indicates a zoomed region in (b) and the white box in (c) is the bottleneck region of interest. The bottleneck problem is observed in the 0.6mm dMRI and 1mm hybrid method as streamlines from the two ROIs are mixed. In the hybrid method at higher resolution (0.6 and 0.4 mm), the streamlines from each ROI demonstrate a separable distribution in the brainstem. d) Using fixel-based analysis, fixels from primary motor cortex (blue), somatosensory cortex (red), and overlapping fixels (pink) are shown (d).

### 3.5 Comparison between hybrid tractography and tracer data

To investigate the neuroanatomical accuracy of hybrid dMRI-PLI tractography, Figure 9 uses a tracer-based connectivity matrix as an estimation of ground truth structural connectivity^11^. The tracer-based connectivity matrix was created by injecting tracers into 29 regions of 91 cortical areas in other macaque brains^12,45,46^. For each injection, the fraction of labelled neurons terminating in each of the 91 areas was calculated, to produce a weighted connectivity matrix of size 29 x 91 (Figure 9B top). To facilitate a fair comparison with the tracer, tractography was also seeded from these 29 injection sites (co-registered to the BigMac brain) and the streamline number projecting to the 91 cortical areas was counted. For each connectivity matrix, the weights were normalized by the total number of connections from each injection site (i.e. the rows sum to one). Figure 9B visually compares the normalized connectivity matrix from the tracer to those generated from dMRI and hybrid dMRI-PLI tractography. As visual comparisons are difficult, we then binarized the tracer matrix using a threshold of 0.0001 (to remove very sparse connections diffusion is unlikely to capture) and generated a receiver operating characteristics (ROC) curve to quantitatively evaluate the sensitivity and specificity of tractography in detecting brain connectivity relative to the tracer (Figure 9C). Here, the dMRI and hybrid matrices were binarized at different thresholds, and the true positive rate (sensitivity) and true negative rate (specificity) with respect to the tracer were calculated. The hybrid method shows superior sensitivity compared to dMRI for the same level of specificity, and superior specificity for the same level of sensitivity, except for when specificity is very high and sensitivity is low where the methods become similar. Figure 9D shows overlap matrices between diffusion/hybrid tractography and tracer for different tractography thresholds. Tracer, tractography, and overlap are visually represented using distinct colours, allowing visualization of false positive, false negative, and true positive connections. At lower thresholds, the hybrid method demonstrates fewer false positive connections compared to dMRI tractography. These results suggest the hybrid method demonstrates neuroanatomical accuracy in estimating brain connectivity above that of typical diffusion tractography.

**Figure 9:**
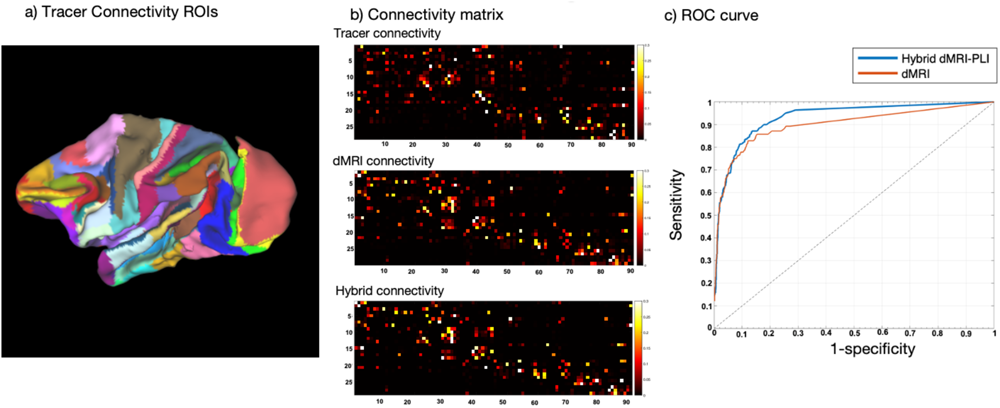
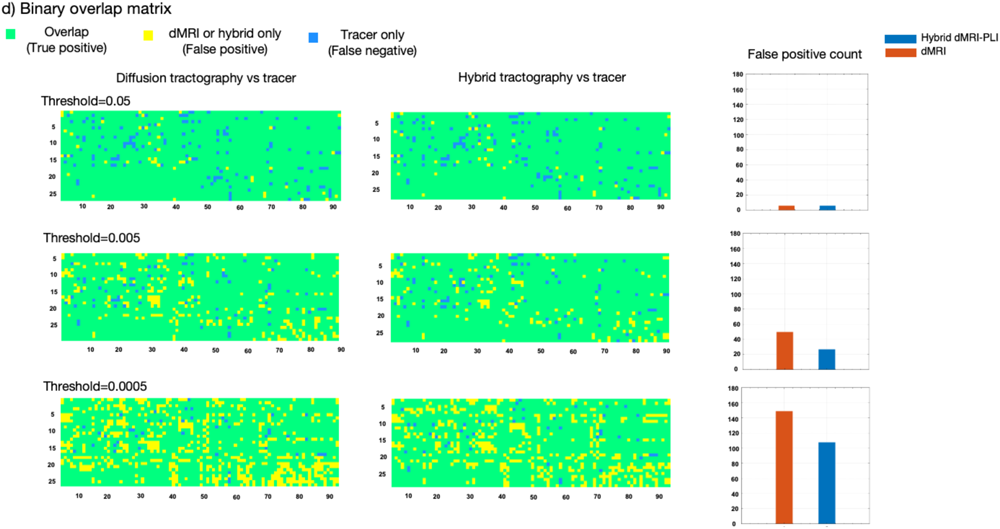
Comparison between the hybrid tractography and dMRI tractography to the tracer connectivity matrix. a) The 91 cortical areas of M132 surface parcellation are shown for the left hemisphere. b) Weighted matrices of size 29 x 91 are shown for the tracer, dMRI tractography, and hybrid tractography. c) An ROC curve illustrating the specificity and sensitivity of hybrid/diffusion-only tractography relative to the tracer connectivity. Different thresholds were applied to the tractography ranging from 0 to 0.25 and the true positive and false positive rates were calculated. d) Binary overlap matrices for different tractography thresholds where tracer only connections are represented in blue (false negative), tractography only in yellow (false positive), and overlap in green (true positive and negative). False positive connectivity was counted for dMRI and hybrid tractography.

### 3.6 Hybrid dMRI-microscopy can be performed with various microscopy contrasts

The hybrid orientations can also be reconstructed from other microscopy contrasts that provide 2D fibre orientation information. The generalisability of the method provides opportunities for other multi-model datasets to reconstruct 3D fibre tractography. Here we demonstrate this using structure tensor outputs from myelin-and Nissl-stained histology also included in the BigMac dataset (Gallyas silver and Cresyl violet stains respectively)^47–50^.

Figure 10 shows how the resulting hybrid FODs at the 0.6mm isotropic resolution are consistent among microscopy contrasts and follow our neuroanatomical expectations. We observe U-shaped fibres, also known as short arcuate fibres, connecting different gyri for three different microscopy contrasts with a consistent curving structure even in the through-plane orientation, the most challenging orientation for the hybrid method. Interestingly, we also observe notable differences in the hybrid outputs from the different microscopy contrasts, where both myelin-and Nissl-stains estimate considerably more crossing-fibre voxels than PLI, as quantified by the crossing fibre ratio calculated across the whole brain. Nonetheless, all three microscopy contrasts facilitate tractography reconstruction of the CST, giving confidence in the generalisability of the method.

**Figure 10:**
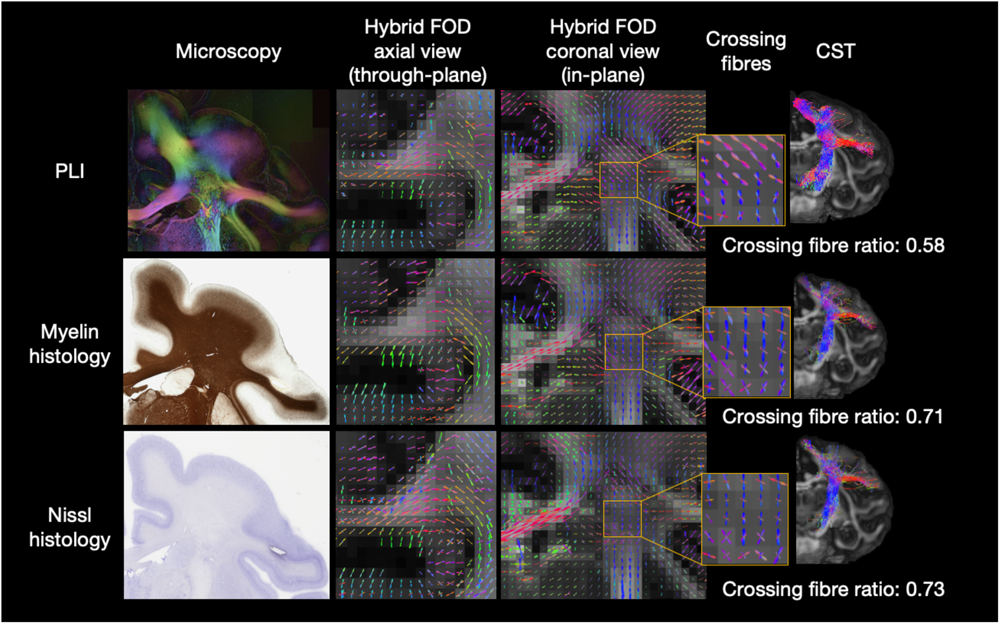
Multiple microscopy contrasts can inform hybrid tractography. Hybrid fibre orientation distributions reconstructed at 0.6mm resolution from different microscopy contrasts: PLI, myelin-and Nissl-stained histology (Gallyas silver/Cresyl violet-stained). All three contrasts show fibre orientations aligned with neuroanatomical expectations. Noticeably, the histology FODs depict more multi-fibre voxels (crossing fibre ratio = *N_wm,mylti-fibre_/N_wm,voxels with fibre_*). These FODs can be fed into tractography to reconstruct white matter tracts (example CST shown).

## 4. Discussion

We develop a method that facilitates microscopy-informed tract reconstruction across the whole brain without relying on invasive tracers. Our method requires data (diffusion MRI and basic light microscopy) that is readily attainable, meaning our method is translatable across species, including humans. Further, white matter tracts can be reconstructed in the same brain allowing study of the microscopy-informed dense connectome (how each cortical region is connected to all other parts of the brain) without having to combine information across subjects. Utilising whole-brain, densely-sampled MRI and microscopy data in the macaque, we construct hybrid dMRI-microscopy fibre orientations that are both informed by high-resolution microscopy and provide 3D information for whole-brain microscopy-informed tractography. We demonstrate how the hybrid orientations can be input into existing tractography algorithms for neuroanatomical investigation of fibre bundles across the brain. Further, we show how the hybrid outputs (FODs and tractography) can be directly compared to dMRI equivalents within the same subject, making the hybrid method a valuable resource for the dMRI modelling community. Notably, the hybrid orientations can be reconstructed at different spatial resolutions using the same underlying data, facilitating meaningful tractography comparisons across different resolutions without SNR confounds that typically complicate similar dMRI-based comparisons. This allows for novel insights into tractography performance, here investigated with respect to the gyral bias and bottleneck problem, that can potentially inspire future algorithms or study designs. In addition, the high-resolution and myelin-specific tractography achievable via the hybrid method can provide neuroanatomical insight, such as interesting patterns of white matter organisation within larger white matter tracts, or reconstruction of the myelinated connectome.

Our data fusion method uses microscopy information to drive tractography-based tract reconstruction with minimum reliance on the dMRI. This is advantageous as it enforces the outputs to be internally consistent with, maximally informed by, and at the high resolution of, the detailed microscopy data. This mindset is considerably different from previous approaches that aim to determine optimal tractography protocols by first performing tractography, comparing these outputs to microscopy/tracing, and using this comparison to update certain parameters or study the influence of various parameters/methods on the tractography sensitivity and accuracy^51–53^. For example, tracts reconstructed from multiple processing pipelines have been compared to tracer masks in the same brain with the aim of determining the most optimal tractography methods^10,51,54^. Alternatively, tractography parameters have been optimised by comparing the tractography-based structural connectome with a tracer-based connectome that combined tracer data across multiple animals to improve brain coverage^55^. More ideally, whole-brain connectome would be performed from 3D microscopy data alone. However, to date this has not been achieved at the whole-brain level in a larger primate brain like the macaque, as 3D microscopy methods are typically restricted to imaging small tissue sections or whole brains from smaller animals (millimetre scale samples)^15,17,20^. This work provides the first whole-brain, microscopy-informed connectome in the macaque, a primate model with complex white matter. Future validation of our method can be performed by comparing our hybrid orientations to 3D PLI^14,30^ acquired on a subset of the microscopy sections.

We investigated whether the hybrid method could address two known challenges in dMRI: the gyral bias and bottleneck problem. Existing dMRI model-based solutions to the gyral bias problem can be limited as they often rely on assumed knowledge of fibre architecture over a gyrus^25,56,57^. This assumption is based on our ability to characterise canonical gyral fanning patterns which are often obtained from invasive imaging methods such as microscopy, involving separate brains or even species. In comparison, the hybrid method directly uses high spatial resolution microscopy data in the same brain. The results (Figure 6) suggest the gyral bias can be reduced and the expected pattern of fibre fanning can be realised with improved spatial resolution^26^. We observe similar trends in both hybrid and diffusion tractography, though the results are more striking in the hybrid method. This may again be related to the fact that the hybrid fibre orientations are more directly related to myelinated axons, where myelin-sensitive microscopy is typically used to characterise fibre fanning patterns ex vivo, and where dendrites complicate dMRI modelling at the grey-white matter boundary. Whether the results can be extrapolated to in vivo data needs further investigation, for example using recently proposed high-resolution dMRI protocols such as Romer-EPTI which is able to acquire sub-mm resolution in an achievable scan time^58^. Considering the global volume difference between humans and macaques (macaque brain volume is ∼1/12 of the human brain volume), the spatial resolutions we used (1mm, 0.6mm and 0.4mm) are approximately equivalent to the 2.5, 1.5, and 1 mm in humans, roughly corresponding to resolutions achievable in the clinic, in typical research data and among the highest achievable resolution so far. Consequently, the resolution examined here in the macaque may provide guidance on solving some tractography challenges in the human brain.

Our hybrid method showed particularly promising results when applied to bottleneck regions, where the hybrid tractography resolved patterns of white matter organisation that mirrored the topographic organisation of the cortex (Figures 7 & <u>8</u>). Bottleneck regions – where multiple fibre bundles converge and share the same orientations-are prevalent in the brain and have been shown to affect a large proportion of white matter tracts leading to false positive streamlines^26,37^. First, we investigated the bottleneck problem in the internal capsule, a relatively thin white matter structure where multiple tracts intermingle to pack within a small volume. Our ability to conserve the order of the motor cortex projections along the anterior-posterior axis in the internal capsule was evaluated. The topographic organisation was conserved in the hybrid tractography at all resolutions including 1mm but lost in the diffusion tractography with 0.6mm. This implies that our ability to resolve the anterior-posterior pattern of white matter organisation in the internal capsule is related to some specific feature of the hybrid FODs, rather than simple improvements in spatial resolution.

For example, the hybrid FODs often have lower dispersion or fewer crossing fibres than the dMRI (likely due to their high specificity to myelin) which may result in streamlines leaving the cortical seed region and following a smooth and continuous fibre bundle instead of jumping between different tracts. Second, we studied the bottleneck region at the brainstem where PLI has been shown to be superior to dMRI when identifying smaller fibre bundles such as lemniscus medialis and pyramidal tracts^32^. Our ability to retain the order of primary motor and somatosensory cortex along the anterior-posterior axis of the brainstem was demonstrated with the hybrid method at higher resolutions (0.6 or 0.4mm), but not in the 1mm hybrid outputs or diffusion tractography at 0.6mm. These results suggest that improved spatial resolution can again help to resolve the bottleneck problem. To translate our findings in vivo, the hybrid method could be used to define masks depicting the separation of streamlines through a bottleneck that could then be utilized in other subjects as tractography waypoint masks to enhance the accuracy of tracking neural pathways through the bottleneck region. Together, hybrid results present a novel way of investigating the bottleneck problem. Our method has the potential to resolve topographic patterns of white matter organisation within major fibre bundles that are currently considered relatively homogeneous or where long-standing hypotheses regarding white-matter topography are currently still lacking empirical evidence^59^.

The hybrid dMRI-PLI tractography is generally well aligned with that of dMRI, though there are also some noticeable differences. To better understand whether the hybrid outputs better reflect ground truth anatomy, we compare the connectivity matrices of dMRI and hybrid tractography to tracer connectivity data from other macaque brains^11^. Our analysis reveals that the hybrid method outperformed dMRI tractography in terms of simultaneously achieving sensitivity and specificity, as indicated by the ROC curve (Figure 9). This suggests that under a lower threshold when false positive connections are a concern, the hybrid method is more effective at detecting true positives and faithfully representing the anatomical connections compared to diffusion tractography. We hypothesise this may be related to the contrast mechanisms driving the FOD reconstruction: the hybrid FOD is based on myelin-sensitive microscopy and so is fairly myelin-specific, whilst the dMRI FOD is influenced by multiple tissue features (e.g. dendrites, glia, and extra-cellular space). Consequently, the hybrid FODs may allow for superior tracking, particularly in regions with considerable partial voluming such as in superficial white matter close to the grey-white matter boundary or when projecting through subcortical structures.

We demonstrate how our method can be applied to different microscopy contrasts, each of which may provide subtly different orientational information that impacts the downstream hybrid tractography (Figure 10). Here, we observe a lower crossing fibre ratio in the hybrid dMRI-PLI FODs, when compared to hybrid FODs reconstructed from either myelin or Nissl-stained histology. This is likely due to the nature of PLI which becomes less informative when a pixel contains multiple fibre populations with different orientations, and the PLI estimation tends to pick up the dominant fibre population^60^. When the two fibre populations are equally weighted and perpendicular to each other, the signals will cancel out. In comparison, the histology data has a finer spatial resolution at 0.28 µm/pixel, where around 200 histology pixels fit into a single PLI pixel (4 µm/pixel), and so can likely better delineate crossing fibre populations or depict fibre dispersion, which may alter or improve the white matter pathways delineation^61^. Consequently, careful consideration should be taken when choosing the microscopy data for hybrid analysis, as different microscopy contrasts provide specificity to different microstructural features. For example, the myelin-stain histology only includes myelinated structures, missing contributions from numerous unmyelinated axons. Nissl-stain in the white matter, on the other hand, visualises glial cell bodies, where oligodendrocytes and astrocyte soma tend to be oriented in short rows running parallel to the surrounding axons^62–64^. Consequently, Nissl-stain provides only an indirect estimate of the fibre orientations. Future work into hybrid outputs from different microscopy contrasts may provide novel neuroanatomical insight into microstructure organisation.

When combining information from MRI and microscopy, we used postmortem MRI rather than in vivo data for several key reasons. First, postmortem MRI provides high SNR data at a higher spatial resolution than is achievable in vivo MRI because postmortem acquisition is less limited by scanning time or motion artefacts caused by physiological processes^65^. Second, the postmortem data provides high angular resolution (i.e. a large number of gradient directions) which has been shown to provide a superior characterisation of crossing fibres and complex fibre architectures^29^. Third, the postmortem MRI is acquired using tissue in the same state as subsequent microscopy (i.e., fixed tissue), negating any microstructural changes due to fixation effects or brain plasticity. Nonetheless, a similar analysis could be conducted using in vivo MRI, assuming the caveats above are considered.

The hybrid dMRI-microscopy method has several limitations. First, it can only be applied to datasets with carefully co-registered MRI and microscopy. We found that tracts reconstructed from the hybrid FODs in the most anterior part of the brain such as the uncinate fasciculus in Figure 4 had less anterior reach compared to the dMRI reconstructions, which in this case more closely followed neuroanatomical expectations. This may be related to errors in the dMRI-microscopy co-registration. dMRI-microscopy co-registration in BigMac was achieved via a state-of-the-art registration tool called TIRL^36^. TIRL provides robust registration of stand-alone microscopy to the volumetric MRI data across most slices. However, due to the limited anatomical information at the most anterior or posterior brain (where sections show little or no white/grey matter contrast), the registration accuracy with these sections is lower. Second, the hybrid method does not provide ground truth information about the tract structure, thus requiring careful consideration when interpreting the neuroanatomical implications of its outputs. Third, the resolution limit of the hybrid FOD is constrained by the density of the microscopy sampling. In BigMac, each microscopy section of the same contrast was repeated every 350 µm, limiting spatial resolution in the anterior-posterior axis to the distance between consecutive slides. Fourth, direct in vivo translation of the hybrid method presented here would be challenging, as it is difficult to obtain microscopy data in vivo. To circumvent these limitations, ongoing work in our lab focuses on using machine learning methods to translate the benefits of the hybrid method in vivo, when only limited dMRI is available.

The hybrid outputs (FODs at different resolutions) are made openly available to the wider community and offer new opportunities for microscopy-informed, whole-brain tractography, where the outputs can be utilised for novel neuroanatomical investigations, or to validate and drive in vivo MRI acquisition and analysis (by helping define the spatial resolution required to observe certain characteristics of the white matter). For example, future work could use the hybrid method to study the superficial white matter short association fibres that connect neighbouring cortical regions with a U-shaped trajectory (“U-fibres”)^66^. These fibres can be challenging to reconstruct using dMRI-data alone due to their highly curved appearance and close proximity to the grey matter^67,68^. However, superficial fibres are often clearly visible within myelin-sensitive microscopy data^69^. Utilising the hybrid methods enhanced sensitivity to myelinated fibre organisation, where the microscopy information acts as an anatomical prior, methods that provide superior sensitivity for tracking U-fibres can be investigated or developed. In addition, fibre tracking within subcortical regions can be performed, such as resolving the hippocampal connectome^70^, thalamic-nuclei segmentation^71^ and the architecture of the cerebellum^72^. Though these regions represent challenging areas for diffusion tractography, the hybrid method based on high-resolution microscopy data, coupled with the high specificity of PLI or myelin-stained histology to axons, may provide superior fibre tracking.

The generalisability of our method opens avenues for hybrid tractography to be performed on other multi-modal datasets that previously could not be used to perform 3D tract reconstruction^13,49^. This includes data acquired from other species facilitating cross-species investigations into structural connectivity, and small tissue blocks that focus on a single white matter bundle or region of interest^41,73^. For a microscopy dataset without dMRI, it is even possible to input dMRI data from another brain or atlas into the model, though the results will be less reliable due to the mismatch of the underlying brain microstructure. Crucially, our method relies on data that can and are being acquired in the human brain (eg. Big Brain project)^74^. Future whole-brain suitable human data will provide exciting opportunities for hybrid reconstruction of the microscopy-informed human connectome, and cross-species comparisons using the macaque data presented here.

## 5. Data and code availability

The pre-processed BigMac data with MRI and microscopy has been previously made openly available via the Digital Brain Bank (https://open.win.ox.ac.uk/DigitalBrainBank/#/). Whole-brain volumes of the fibre orientation distributions output from the hybrid method will be made openly available. This includes dMRI-PLI FODs at multiple spatial resolutions (1, 0.6 and 0.4mm isotropic), hybrid FODs from myelin-and Nissl-stained histology at 0.6mm, and dMRI FODs at 0.6mm. The code for constructing the hybrid orientation from dMRI and microscopy is accessible via (https://git.fmrib.ox.ac.uk/srq306/hybrid_microscopy_dmri) with further tutorial documentation provided at (https://open.win.ox.ac.uk/pages/srq306/hybridtractographydoc/).

## 6. Methods

### 6.1 MRI data and microscopy data acquisition

The BigMac dataset was previously acquired and pre-processed as described in Howard et al., 2023^29^. Relevant to this work, an adult rhesus macaque brain was scanned postmortem on a 7T small animal scanner. Structural images were acquired with multi gradient echo sequence at a spatial resolution of 0.3mm isotropic: FOV =76.8 × 76.8 × 76.8 mm, TE/TR = 7.8/97.7 ms, and flip angle = 30°.

Postmortem dMRI was acquired with spin echo 2D multi-slice sequence and single-line readout: 0.6mm, 128 gradient directions at b=4ms/µm^2^ and 8 with negligible diffusion weighting.

After the scanning, the brain was sectioned into two blocks (anterior/posterior halves). Each block was sectioned into thin (50/100 µm) slices and allocated to one of six interleaved contrasts: polarised light imaging (PLI)^16,30–32^, Cresyl violet staining for Nissl bodies^34,35^, Gallyas silver staining for myelin^33^ and three unassigned sections that were stored for longevity. The slice thickness was 50 *μm* for 5 out of the 6 sections (including PLI, Nissl and myelin-stained sections) with one section 100 *μm* thick. Each section of the same contrast was repeated every 350 µm.

PLI estimated the primary fibre orientation based on the birefringence of myelinated axons with a resolution of 4 µm per pixel^16,30–32^. Maps of fibre orientation within the microscopy plane (in-plane angles) were used as input for the hybrid dMRI-PLI tractography.

Histology slides with Gallyas silver staining (myelin) and Cresyl violet staining (Nissl bodies) were digitised at a spatial resolution of 0.28 µm/pixel. 2D structure tensor analysis^47–49^ of the stained sections (using a Gaussian kernel with sigma=10 pixels) was used to estimate the fibre orientations for each microscopy pixel.

### 6.2 MRI-Microscopy data co-registration

MRI-Microscopy registration was performed using TIRL (v3.1.1), an automated pipeline facilitating the accurate registration of the histology sections to the whole brain MRI via a sequence of linear and non-linear transformations (see ^29,36,75,76^ for further details). Here each PLI, myelin-and Nissl-stained slide was registered independently to the MRI space. The structural MRI was used as a reference due to its high resolution and good grey/white contrast. Separately, the structural and dMRI were co-registered using linear registration in FSL (FLIRT)^77,78^. By combining the TIRL and FLIRT transformations, the microscopy pixels were mapped to voxel coordinates in dMRI space. Mapping of the microscopy fibre orientations (vectors) to dMRI space also accounted for local rotations according to the warp field.

### 6.3 Diffusion-only orientation estimation

Fibre orientations were estimated from dMRI for two purposes. First, we required orientation estimates to be input into the hybrid method and provide the through-plane orientation information. Here, the dMRI was analysed using the ball and stick (BAS) model^27^ to estimate ≤ 3 fibre populations with 50 orientation estimates (samples) per population. The BAS model was chosen due to its ability to estimate multiple fibre populations per voxel and discrete orientation samples that can be directly compared with those from microscopy. Other diffusion models providing estimation for the fibre orientation could also be used.

Second, we analysed the data using a spherical-harmonic-based method, constrained spherical deconvolution (CSD)^79^, where the CSD-derived FODs and downstream tractography could be directly compared to equivalent outputs from the hybrid method (where our FODs are similarly described using spherical harmonics). Here, CSD was performed using the dhollander algorithm for fibre response function estimation^80,81^.

### 6.4 Hybrid orientation estimation

To calculate the 3D hybrid dMRI-microscopy orientations, microscopy provided the fibre orientations within the microscopy plane, and we approximated the through-plane angle (or inclination angle) with that from dMRI (BAS model). A schematic of the method is illustrated in Figure 1 with additional details provided in <u>Supplementary</u> Figure 3.

PLI and myelin-/Nissl-stained structure tensor analysis provided an estimate of the in-plane orientation per microscopy pixel. The in-plane orientation was warped to dMRI space and compared with the co-registered BAS samples that were projected onto the microscopy plane. Samples from BAS populations with signal fractions less than 0.05 were excluded. These sample orientations captured the uncertainty in the estimation of different fibre populations. The through-plane angle was then approximated with that from the most similar BAS sample to produce a 3D hybrid orientation per microscopy pixel. Thus, we assigned the through-plane orientation of the BAS samples to their precise sub-voxel localisation and informed the 3D configuration of fibre orientations from the microscopy.

Next, orientations were combined over a local neighbourhood into a fibre orientation distribution (FOD) defined by a frequency histogram of fibre orientations for 256 evenly spaced points across a sphere. Spherical harmonics of order 8 were fitted to the histogram. FODs were reconstructed at the spatial resolution of 1.0, 0.6, and 0.4 mm isotropic. dMRI-derived (CSD) FODs are typically not normalised for the integral to sum to one. To ensure our hybrid FODs had amplitudes similar to those from dMRI (i.e. to make the tractography outputs comparable), the hybrid FODs were scaled on a voxel-wise basis so that the amplitude of the maximum hybrid/dMRI-CSD FOD peak was equivalent. This scaling method was chosen based on the visual evaluation of comparable outputs from the hybrid and diffusion tractography.

In BigMac, there was a region in the centre of the brain (approx. 1.8-3 mm thick) with missing microscopy because the brain was sectioned in two halves (an anterior and posterior block). In this region, the missing data were filled with the FODs from the dMRI data estimated via CSD^79^ (<u>Supplementary</u> Figure 5). To fill the missing data of the hybrid orientation at 0.4mm and 1mm, the CSD FODs were up-/down-sampled as appropriate with MRtrix3^82^.

### 6.5 Tractography

Tractography was performed using MRtrix3^82^ and FODs either derived from the hybrid method above (hybrid tractography), or from CSD^81^ of the dMRI data (diffusion tractography). Global fibre tractography was achieved using the iFOD2 algorithm^38^ with the following parameters: 30 seeds per voxel in the white matter, step size of 0.2, cut-off value of 0.05, maximal length of 120mm, and minimal length of 5mm. Fibre bundle segmentation was performed with inclusion and exclusion masks from XTRACT to extract 42 tracts spanning the whole brain, including association, commissural, limbic and projection fibres^40^. The masks were warped from standard (F99) space to the BigMac diffusion space.

The reconstructed tracts were further refined to remove the spurious streamlines (<u>Supplementary</u> Figure 3). First, each streamline within a given tract was compared to the tract probability map from the population-based XTRACT white matter atlas obtained from 6 macaque brains^40^. The streamline was assigned a score based on the mean probability value of the voxels the streamline intersected. If the mean probability of a streamline was lower than 0.2, it was discarded^83^. Second, streamlines that were 4 standard deviations longer than the mean fibre length or deviated over 5 standard deviations relative to the core of the fibre were removed. The parameters were chosen based on the literature and were fine tuned for our data^83^. The density map of each tract was calculated on a voxel-wise basis as the number of streamlines per voxel divided by the total number of streamlines of the tract.

### 6.6 Gyral bias

Gyral crowns are defined as regions of high convexity and gyral walls are defined by their low curvature. FODs near the gyri and gyral streamline structure were examined for different spatial resolutions (0.4, 0.6 and 1mm). Global tractography was then performed with the parameters mentioned above, but now with 5 seeds per voxel to facilitate easier visualisation of streamlines near the cortex.

### 6.7 Bottleneck regions

We investigated two bottleneck regions within the motor network to determine whether hybrid tractography with different spatial resolutions can retain the topographic organization of fibre bundles when passing through the bottleneck regions. First, we investigated whether the topographic medial-lateral representation of the trunk, arms and face areas across the motor cortex would be retained for fibres projecting through the internal capsule, as has been previously described^42^. Second, we studied fibre bundles projecting from the primary motor and somatosensory motor cortex into the brainstem to determine whether the anterior-posterior distribution was retained^44^.

The ROIs in the motor/somatosensory cortex were manually drawn on the structural MRI surface (Figure 7 and Figure 8) and then converted into volume space, warped to the dMRI and used for tractography seed masks. The internal capsule mask was obtained from the D99 atlas v2.0^84^ and the brainstem mask was obtained from the subcortical atlas of the Rhesus Macaque (SARM)^85^. Both were warped to BigMac diffusion space. Seeding from the ROIs in the cortex, tractography was performed following the same parameters as above. Only streamlines going through the internal capsule or brainstem were retained. Tract density maps were generated for each ROI. The masks within the internal capsule and brainstem were colour-coded based on the ROI with the highest number of streamlines. The outputs in the internal capsule and brainstem were compared for both hybrid orientations and dMRI data.

To identify the bottleneck area, fixel-based analysis was performed with MRtrix3^82^. A fixel describes a specific fibre population within a voxel. Tracts generated from each seed ROI were converted to fixel density maps using the tck2fixel command. Here the FOD was first segmented into discrete fixels and each streamline passing through a voxel was assigned to the fixel that exhibited best alignment^43^. The fixel density map was then thresholded at 5% and converted into a fixel mask. The bottleneck area was identified as the region where multiple fibre bundles use the same fixel when fibre tracking through the voxel (“multi-bundle fixels”).

### 6.8 Comparison with the tracer tracking

We compared our hybrid tractography to a weighted connectivity matrix derived from retrograde tracer data, which serves as a ground truth estimate of structural connectivity^11^. We used a weighted connectivity matrix constructed from tracer injections in 28 monkeys to examine the connectivity profiles of cortical-cortical pathways. Tracers were injected into 29 regions of 91 cortical areas from the macaque atlas in the left hemisphere^12,45,46^, and the fraction of labeled neurons (FLNe), defined as the number of labeled neurons in each cortical area relative to the total number of neurons in the injection, was used to quantify connectivity. As we aimed to investigate the improvement of hybrid tractography over dMRI tractography with the tracer connectivity as a reference, we applied a threshold of 0.0001 to exclude very sparse tracer connections that are unlikely to be captured by tractography^11^. The cortical atlas was co-registered to BigMac space and hybrid/diffusion-only tractography was seeded from the 29 injection sites of the left hemisphere with the tractography parameters mentioned above. The number of streamlines projecting to each of the 91 cortical regions, divided by the total number of streamlines from that injection site, was used to construct the tractography connectivity matrix (29 x 91 areas).

To facilitate quantitative comparison via a ROC curve, the tracer matrix was first binarized using a threshold of 0.0001. The hybrid/diffusion-only connectivity matrices were then also binarized using thresholds ranging from 0 to 0.25 and the number of true positives (TP), false positives (FP), true negatives (TN) and false negatives (FN) counted for each threshold. The sensitivity was calculated as the true positive rate TPR=TP/(TP+FN) and specificity as the true negative rate TNR=TN/(TN+FP). The ROC curve was then plotted as sensitivity versus 1-specificity. Overlap matrices (hybrid/diffusion tractography versus tracer) were then plotted for different tractography thresholds with the tracer in yellow, tractography in blue, and overlap in green.

## 8. Contributions

All authors reviewed the manuscript. SZ: project lead; involved with hybrid dMRI-microscopy pipeline development, conceived of and performed analyses; wrote and edited manuscript. INH: developed the TIRL software and protocol for co-registration of the MRI-microscopy data. MC, GD, TH, AK, RM, JM, CS, AS etc were involved in the acquisition and curation of the BigMac data. NE: advised on the cortical ROI drawing. SJ: study conceptualisation. KLM: study conceptualisation, advised on all processing pipelines and analyses; evaluated the results, edited manuscript; provided resources, supervision and funding. AFDH: study conceptualisation, involved with hybrid dMRI-microscopy pipeline development, advised on all processing pipelines and analyses; evaluated the results, edited manuscript.

## 6. Supplementary figures

**Supplementary figure 1:**
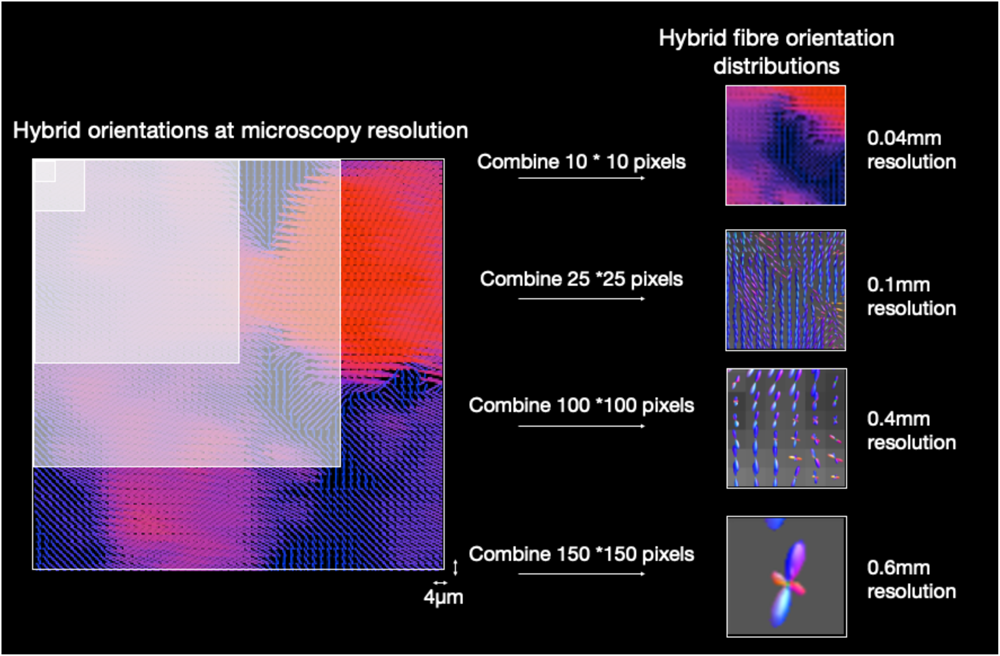
Reconstruction of hybrid fibre orientation distributions at various resolutions. The hybrid method generates 3D fibre orientations at the resolution of the microscopy data that can then be combined over a local neighbourhood into a fibre orientation distribution (FOD). The resolution of the hybrid FOD depends on the size of the combined region. Using 4μm resolution microscopy, here we demonstrate how a 0.04mm FOD can be generated by combining 10*10 pixels, 0.1mm with 25*25 pixels, 0.4mm with 100*100 pixels or 0.6mm with 150* 150 pixels. For simplicity, the example is shown in 2D but our approach was implemented in 3D (see Supplementary Figure 2).

**Supplementary figure 2:**
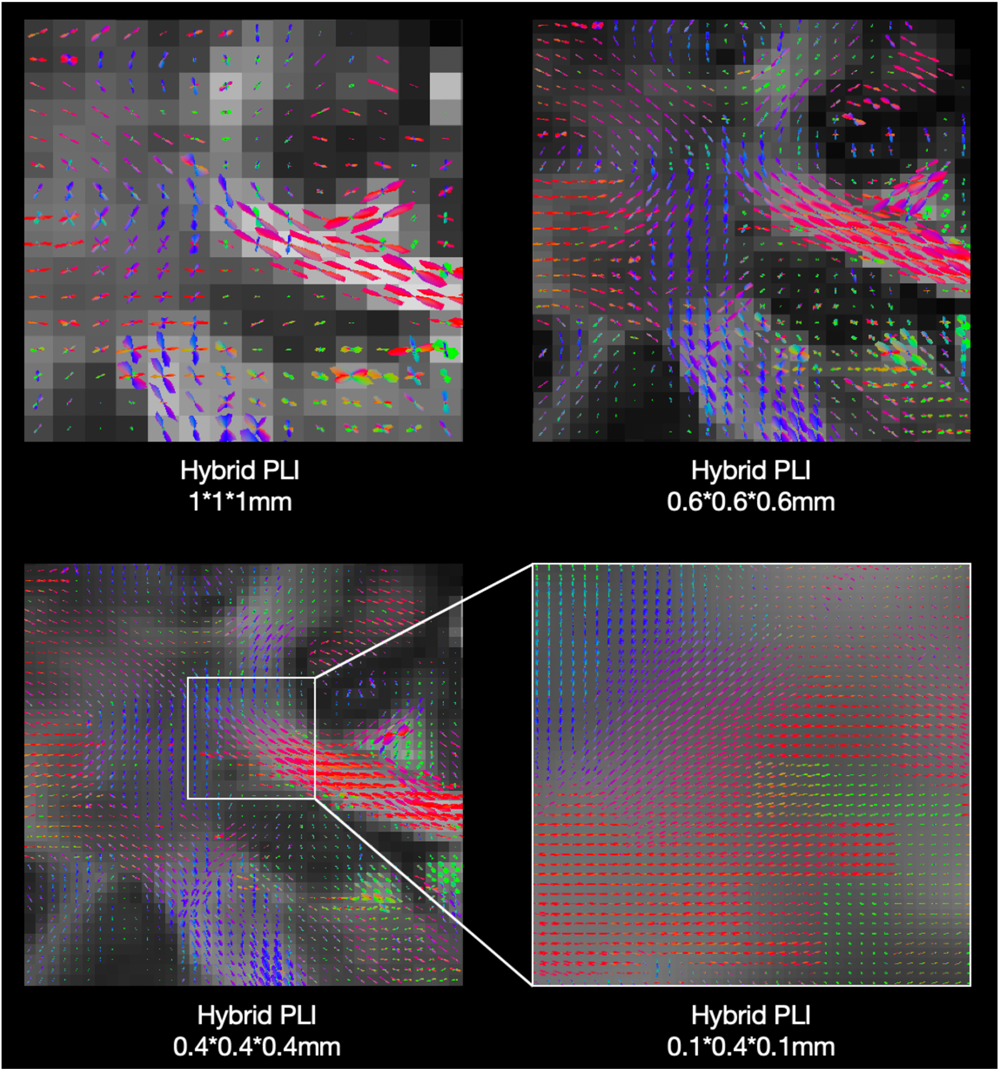
Hybrid fibre orientation distributions reconstructed at different spatial resolutions. Here we show example outputs for 1*1*1mm, 0.6*0.6*0.6mm, 0.4*0.4*0.4mm and 0.1*0.4*0.1mm (XYZ). X-Z denotes the coronal plane where microscopy was acquired with a native resolution of 4 μm/pixel, whilst Y denotes the axis along which the brain was sectioned, where microscopy sections of similar contrast are repeated every 0.35mm throughout the brain. Hence the asymmetric 0.1*0.4*0.1mm voxels. The fibre orientations vary smoothly at all resolutions.

**Supplementary figure 3:**
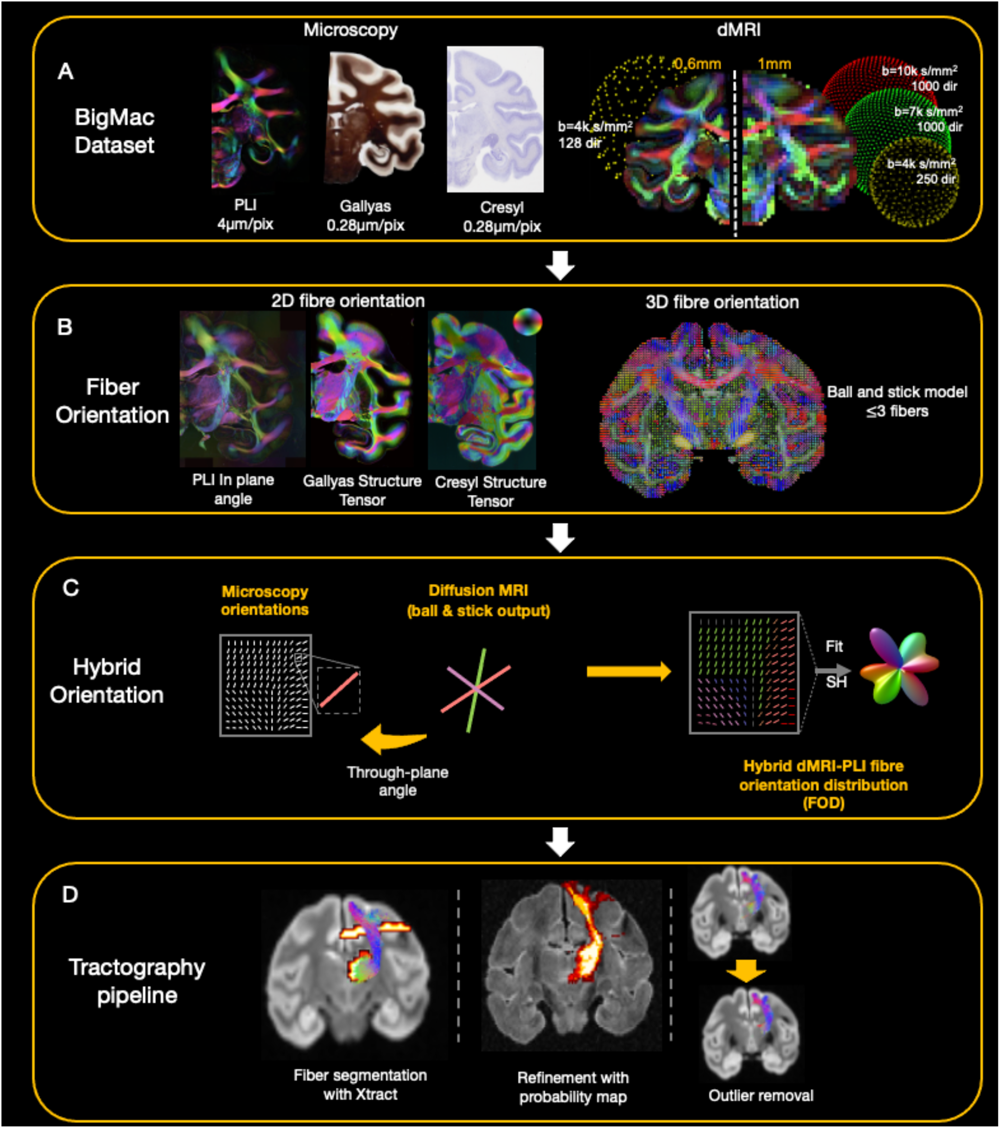
The overview of the analysis pipeline. A) BigMac includes co-registered microscopy (PLI, myelin-and Nissl-staining) and postmortem MRI (dMRI/structural). B) Fibre orientations are extracted from each microscopy contrast (in-plane angle from PLI, structure tensor analysis from histology) and dMRI (ball and stick model). C) Hybrid orientations are generated with the in-plane orientation from microscopy and through-plane orientation from dMRI. D) The tractography results are optimized with XTRACT probability maps and outlier removal.

**Supplementary figure 4:**
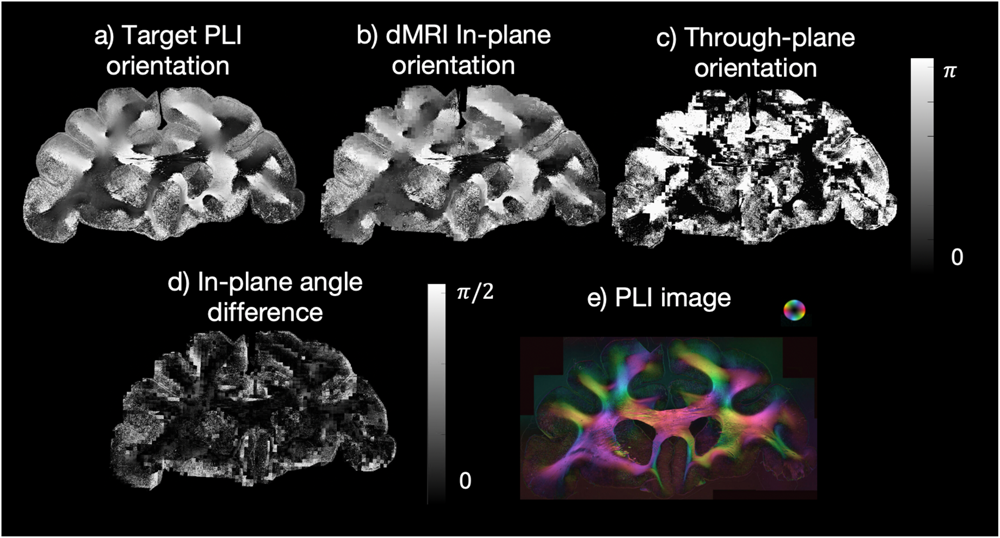
The angle component used in the hybrid orientation reconstruction. a) The PLI orientation; b) The in-plane orientation from the diffusion BAS decomposition (represented by the angle projected onto the microscopy plane); c) The through-plane orientation from the diffusion BAS decomposition (inclination angle); d) The in-plane angle difference between the target PLI orientation and the most similar BAS fibre orientation which has been projected onto the microscopy plane showing a small angle difference. e) The PLI hue-saturation-value with the colour-coded orientations.

**Supplementary figure 5:**
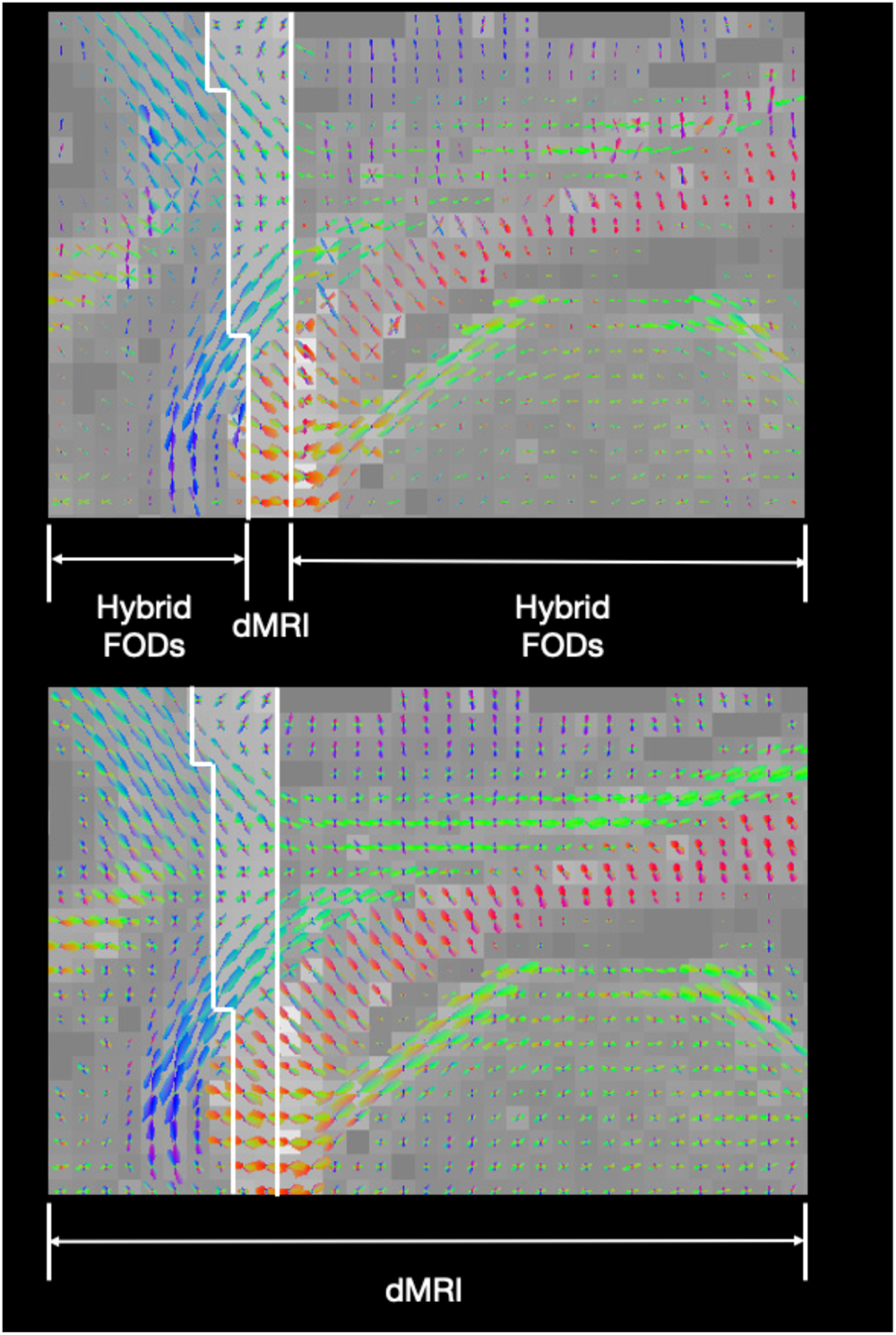
Missing microscopy filled with dMRI CSD. The BigMac brain was sectioned into two halves (anterior & posterior), resulting in a gap of approximately 3-5 voxels without microscopy data in the centre of the brain. To facilitate whole-brain tractography using the hybrid method, these voxels were filled by FODs from dMRI-CSD. The continuity and similarity of the FODs after filling (Top) is shown in comparison with dMRI CSD (bottom).

**Supplementary figure 6:**
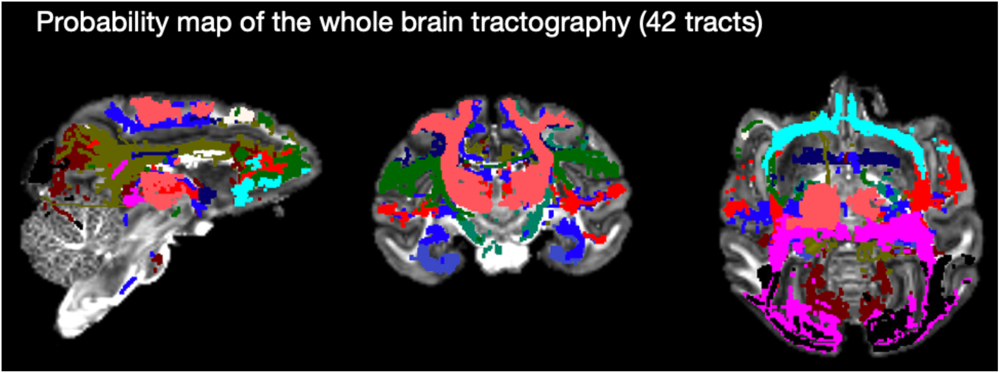
Tracts spanning the whole brain reconstructed using hybrid dMRI-PLI at 0.6mm. XTRACT was used to reconstruct 42 major white matter tracts using the hybrid FODs. Here we overlay tract masks to demonstrate the substantial coverage of the 42 tracts across the brain.

**Supplementary figure 7:**
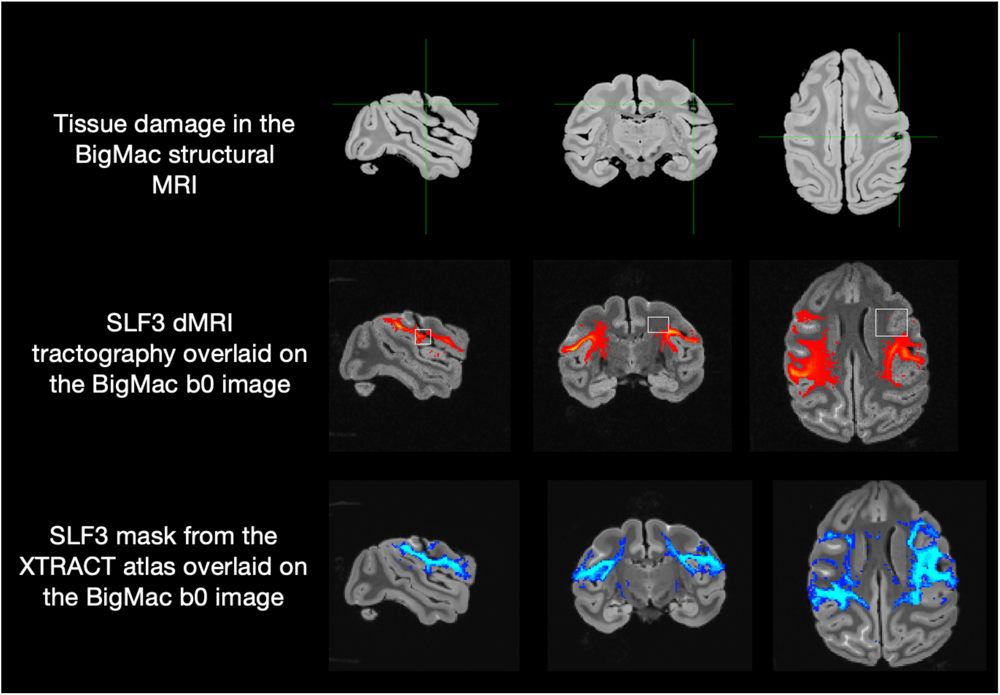
Bleeding site in the BigMac dataset. The tissue damage in the left hemisphere is evident in the postmortem MRI structural image (Top). This affected the tracking of the superior longitudinal fasciculus 3 (SLF3) (middle) compared to the XTRACT atlas (bottom), with missing regions indicated by the white boxes.

**Supplementary figure 8:**
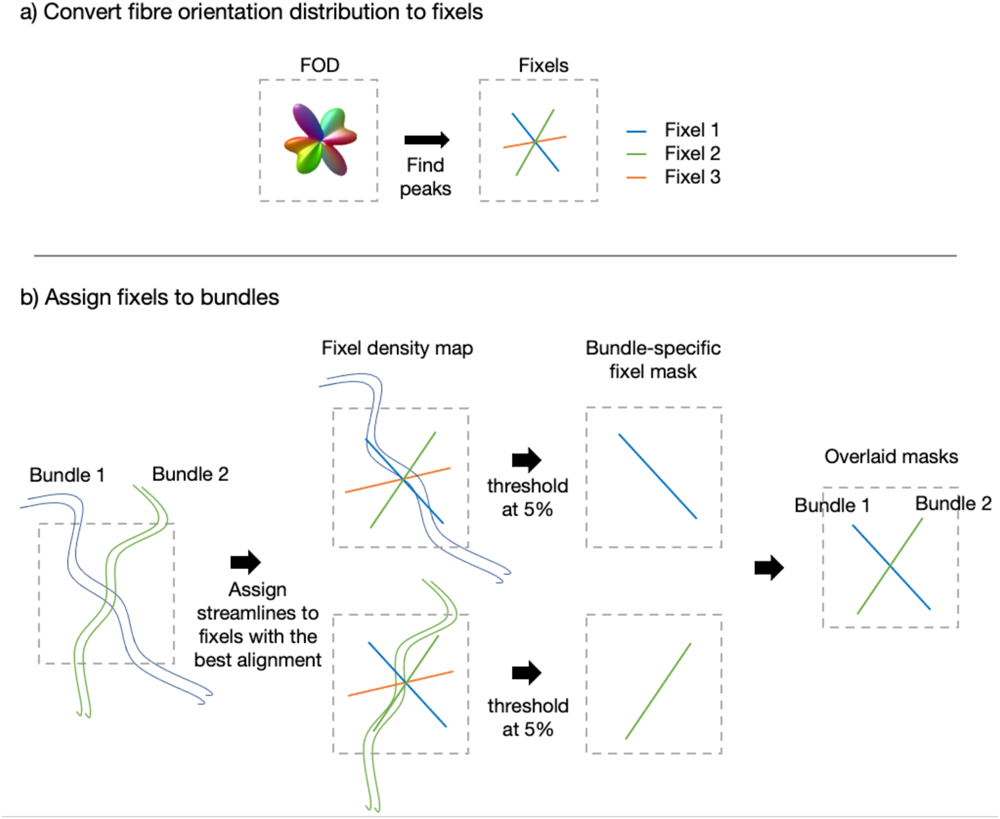
Fixel-based analysis. a) The fibre orientation distribution can be subdivided into “fixels”, each of which describes a specific fibre population within the voxel. b) Each streamline is assigned to a specific fixel. This allows us to compute a fixel density map for each fibre bundle. The fixel density map is calculated for each fibre bundle independently and describes the number of streamlines from a given fibre bundle that passes through each fixel. The density map is converted into a fixel mask by setting a threshold (here 5%). We can then combine the masks across fibre bundles to visualise whether they pass through the voxel with different orientations (as shown) or the same fibre orientations (indicating a bottleneck). The tck2fixel in MRtrix3^46^ was utilized to implement this procedure.

